# Longitudinal Metaproteomic and Lysine Crotonylation Remodeling Defines Clinical Response to Fecal Microbiota Transplantation in Ulcerative Colitis

**DOI:** 10.64898/2026.10.13.758943

**Authors:** Yangcheng Liu, Wanning Chen, Yue Yuan, Xin Liu, Zhangqin Li, Bo Li, Lixin Zhu, Ruixin Zhu, Yinglei Miao, Weili Lin, Yang Sun

**Affiliations:** Department of Gastroenterology, The First Affiliated Hospital of Kunming Medical University, Yunnan, P.R. China; Shanghai Key Laboratory of Maternal Fetal Medicine, Shanghai Institute of Maternal-Fetal Medicine and Gynecologic Oncology, Clinical and Translation Research Center, Shanghai First Maternity and Infant Hospital, School of Life Sciences and Technology, Tongji University, Shanghai 200092, P.R. China; Institutes of Biomedical Sciences, School of Life Sciences, Inner Mongolia University, Hohhot 010070, P.R. China

**Keywords:** Fecal microbiota transplantation, ulcerative colitis, clinical response, metaproteomics, lysine crotonylation, post-translational modification, microbiome

## Abstract

**Background:** Fecal microbiota transplantation (FMT) can induce steroid-free clinical remission in a subset of patients with ulcerative colitis (UC), but treatment responses remain heterogeneous and are not fully explained by taxonomic microbiome shifts. To define functional molecular features associated with FMT efficacy, we performed integrated longitudinal fecal metaproteomic and lysine crotonylation (Kcr) profiling.

**Methods:** We analyzed 87 fecal samples from 23 patients with active UC collected at baseline and at weeks 1, 8, and 12 after FMT. Patients were classified as responders (n = 13) or non-responders (n = 10) according to steroid-free clinical remission at week 12. Across samples, we identified an average of 3,682 proteins and 562 Kcr sites per sample.

**Results:** Responders showed greater metaproteomic and meta-crotonylomic complexity at week 12, sustained enrichment of *Prevotella* and *Faecalibacterium*, and eight directionally consistent baseline-altered proteins with candidate prognostic value. After FMT, responders exhibited coordinated microbial functional remodeling, including increased proteins involved in short-chain fatty acid (SCFA) metabolism, acyl-CoA-centered pathways, nutrient transport, and polysaccharide degradation, together with attenuation of host neutrophil-related inflammatory protein and Kcr signatures. Non-responders showed minimal or opposing molecular changes.

**Conclusions:** These findings nominate longitudinal fecal metaproteomic and Kcr remodeling as functional molecular readouts of clinical response to FMT in UC and highlight lysine crotonylation as an underexplored metabolite-sensitive layer linking microbial metabolic restoration with host inflammatory resolution.

**Importance:** Fecal microbiota transplantation can send ulcerative colitis into remission, yet clinicians have no reliable way to predict who will respond. By tracking 23 patients for 12 weeks, researchers show that stool proteins and a chemical mark called lysine crotonylation reveal the answer. Future responders carried distinct baseline protein signatures, and after treatment their gut microbes restarted key metabolic engines—short-chain fatty acid and acyl-CoA pathways—while host neutrophil inflammation quieted down. Non-responders lacked this coordinated rebound. The work delivers a functional molecular toolkit to forecast and monitor transplant success, shifting the focus from cataloging gut bacteria to decoding the real-time dialogue between microbial restoration and immune calm.

## INTRODUCTION

Ulcerative colitis (UC) is a chronic relapsing inflammatory bowel disease (IBD) characterized by mucosal inflammation and substantial inter-individual variability in therapeutic response^1-3^. Although fecal microbiota transplantation (FMT) has emerged as a clinically relevant microbiota-targeted intervention for UC^4-6^, only a subset of patients achieve durable clinical and endoscopic remission. Current assessment of FMT efficacy relies primarily on clinical indices and endoscopy^7^, whereas the functional molecular processes that distinguish responders from non-responders remain insufficiently defined. A key translational gap is the lack of longitudinal molecular readouts that capture therapeutic response beyond taxonomic community shifts.

Metaproteomics directly profiles functionally active microbial and host proteins in the intestinal environment and can capture biological processes that are not apparent from DNA-based profiling alone^8^. However, protein abundance does not fully represent the post-translational regulatory layer of host-microbiome interactions. Recent work has shown that microbial protein acylation is widespread in the gut microbiome and altered in IBD^10^. Lysine crotonylation (Kcr) is a metabolite-sensitive acylation modification linked to intracellular acyl-CoA availability and microbial short-chain fatty acid (SCFA) metabolism^11,12^. Although crotonylation has been implicated in mammalian inflammatory and epithelial programs, whether Kcr occurs on gut microbial proteins in vivo and how fecal Kcr remodeling relates to clinical FMT response in UC remain largely unexplored.

Here, we integrated longitudinal fecal metaproteomics with Kcr profiling in patients with active UC undergoing FMT. By following patients from baseline through weeks 1, 8, and 12 after treatment, we aimed to identify candidate pretreatment molecular features, define treatment-emergent protein and Kcr remodeling, and link microbial metabolic restoration with host inflammatory resolution. This design positions Kcr as a candidate functional layer for monitoring clinical FMT response beyond conventional taxonomic profiling.

## RESULTS

### Clinical characteristics and outcomes of UC patients undergoing FMT

A total of 23 patients with active UC who underwent FMT and provided longitudinal stool samples were included (Fig. 1A). Patients were classified as responders (n = 13) or non-responders (n = 10) according to steroid-free clinical remission at week 12, defined as a total Mayo score ≤ 2 and an endoscopic Mayo subscore ≤ 1. Baseline clinical and demographic characteristics were balanced between groups (Table 1). In total, 87 fecal samples were available for integrated fecal metaproteomic and Kcr analysis across four time points: baseline (responders, n = 11; non-responders, n = 10), week 1 (responders, n = 13; non-responders, n = 10), week 8 (responders, n = 13; non-responders, n = 10), and week 12 after FMT (responders, n = 12; non-responders, n = 8).

**Table 1.**
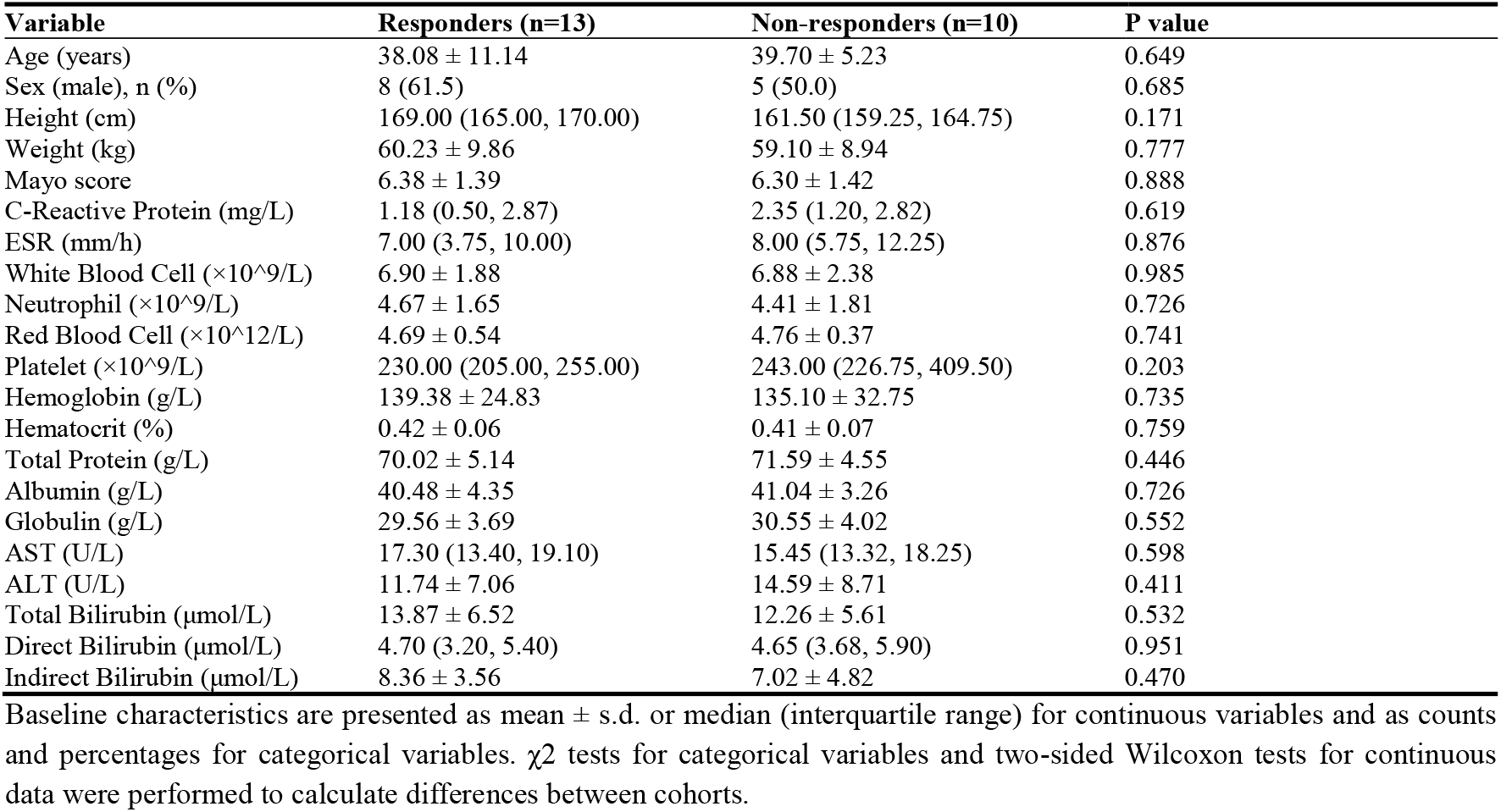
Baseline characteristics of UC patients.

| Variable | Responders (n=13) | Non-responders (n=10) | P value |
| --- | --- | --- | --- |
| Age (years) | 38.08 ± 11.14 | 39.70 ± 5.23 | 0.649 |
| Sex (male), n (%) | 8 (61.5) | 5 (50.0) | 0.685 |
| Height (cm) | 169.00 (165.00, 170.00) | 161.50 (159.25, 164.75) | 0.171 |
| Weight (kg) | 60.23 ± 9.86 | 59.10 ± 8.94 | 0.777 |
| Mayo score | 6.38 ± 1.39 | 6.30 ± 1.42 | 0.888 |
| C-Reactive Protein (mg/L) | 1.18 (0.50, 2.87) | 2.35 (1.20, 2.82) | 0.619 |
| ESR (mm/h) | 7.00 (3.75, 10.00) | 8.00 (5.75, 12.25) | 0.876 |
| White Blood Cell ($\times 10^9/L$) | 6.90 ± 1.88 | 6.88 ± 2.38 | 0.985 |
| Neutrophil ($\times 10^9/L$) | 4.67 ± 1.65 | 4.41 ± 1.81 | 0.726 |
| Red Blood Cell ($\times 10^{12}/L$) | 4.69 ± 0.54 | 4.76 ± 0.37 | 0.741 |
| Platelet ($\times 10^9/L$) | 230.00 (205.00, 255.00) | 243.00 (226.75, 409.50) | 0.203 |
| Hemoglobin (g/L) | 139.38 ± 24.83 | 135.10 ± 32.75 | 0.735 |
| Hematocrit (%) | 0.42 ± 0.06 | 0.41 ± 0.07 | 0.759 |
| Total Protein (g/L) | 70.02 ± 5.14 | 71.59 ± 4.55 | 0.446 |
| Albumin (g/L) | 40.48 ± 4.35 | 41.04 ± 3.26 | 0.726 |
| Globulin (g/L) | 29.56 ± 3.69 | 30.55 ± 4.02 | 0.552 |
| AST (U/L) | 17.30 (13.40, 19.10) | 15.45 (13.32, 18.25) | 0.598 |
| ALT (U/L) | 11.74 ± 7.06 | 14.59 ± 8.71 | 0.411 |
| Total Bilirubin ($\mu\text{mol/L}$) | 13.87 ± 6.52 | 12.26 ± 5.61 | 0.532 |
| Direct Bilirubin ($\mu\text{mol/L}$) | 4.70 (3.20, 5.40) | 4.65 (3.68, 5.90) | 0.951 |
| Indirect Bilirubin ($\mu\text{mol/L}$) | 8.36 ± 3.56 | 7.02 ± 4.82 | 0.470 |

Baseline characteristics are presented as mean ± s.d. or median (interquartile range) for continuous variables and as counts and percentages for categorical variables. $\chi^2$ tests for categorical variables and two-sided Wilcoxon tests for continuous data were performed to calculate differences between cohorts.

**Fig. 1.**
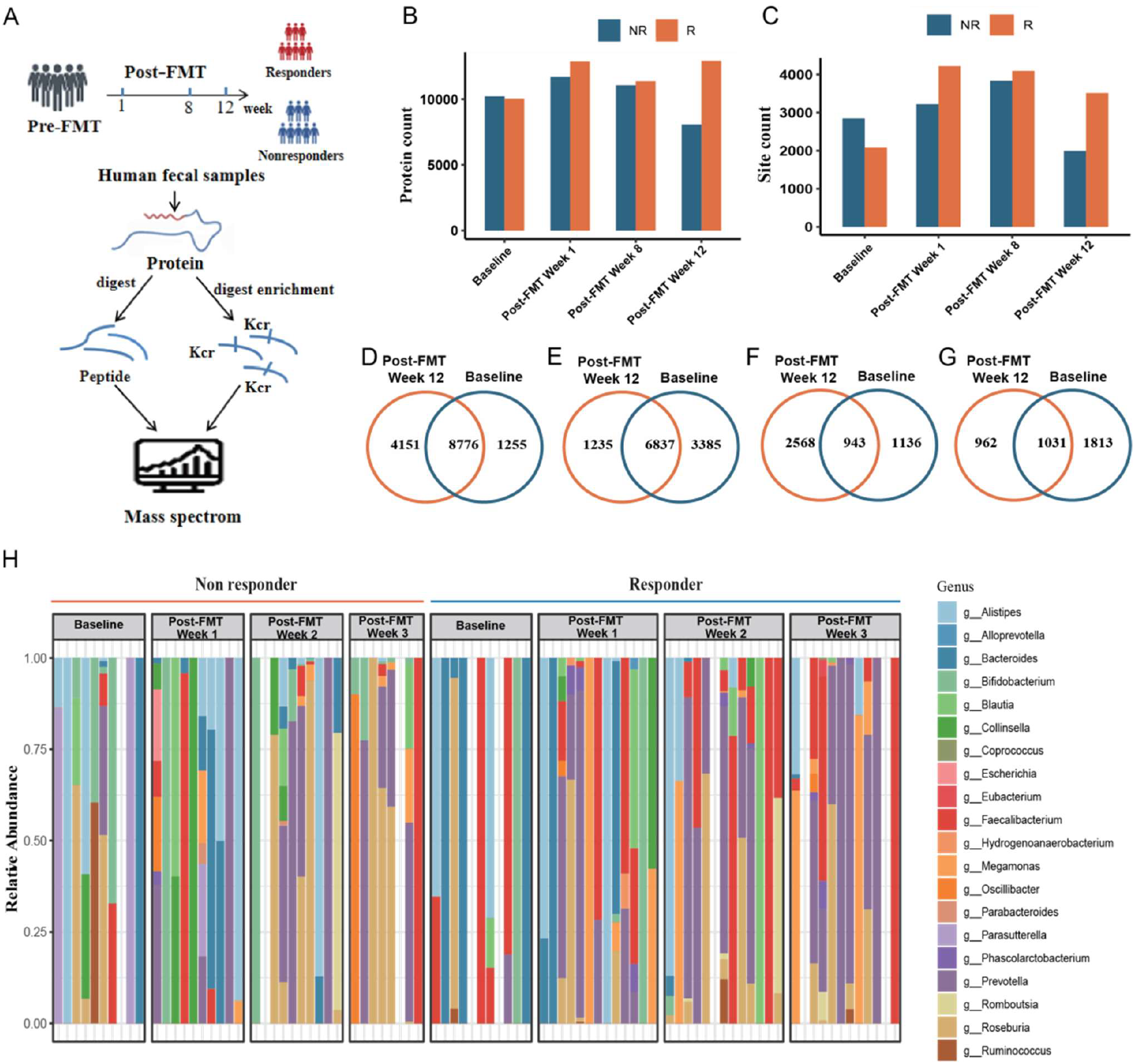
Experimental workflows and overview of metaproteomic and crotonylation profiles. (A) Schematic of integrated sample processing for metaproteomics and Kcr enrichment. (B, C) Bar plots showing counts of identified proteins (B) and Kcr sites (C) at each time point in responders (R) and non-responders (NR). (D-G) Venn diagrams illustrating overlap of proteins (D, E) and Kcr sites (F, G) between baseline and week 12. (H) Heatmap showing genus-level relative abundance of taxa based on quantified peptides.

### Longitudinal metaproteomic and meta-crotonylomic profiles capture response-associated molecular complexity

We performed global metaproteomic and Kcr profiling using an integrated enrichment strategy (Fig. 1A). Across all 87 samples, we identified an average of 3,682 proteins and 562 Kcr sites per sample. At week 12 after FMT, responders exhibited significantly higher numbers of detected proteins and Kcr sites than non-responders (Figs. 1B, 1C), consistent with greater response-associated functional and post-translational molecular complexity. Temporal overlap analysis further showed that Kcr sites underwent substantially greater turnover than total protein abundance after FMT, with only approximately 45% and 36% of baseline Kcr sites retained at week 12 in responders and non-responders, respectively (Figs. 1D-1G). Taxonomic inference from quantified peptides showed that responders maintained higher relative abundances of Prevotella and Faecalibacterium throughout treatment (Fig. 1H).

### Baseline and treatment-emergent protein/Kcr signatures distinguish FMT responders from non-responders

Longitudinal comparisons revealed marked molecular divergence between responders and non-responders. At baseline, 182 microbial proteins and 12 microbial Kcr sites differed significantly between groups (Figs. 2A, 2E), indicating pre-existing functional differences between patients who subsequently did or did not achieve remission. These differences persisted and evolved after FMT, with differential features remaining evident at weeks 1, 8, and 12 (Figs. 2B-2D, 2F-2H). Differential human protein and Kcr features are summarized in Fig. S1.

**Fig. 2.**
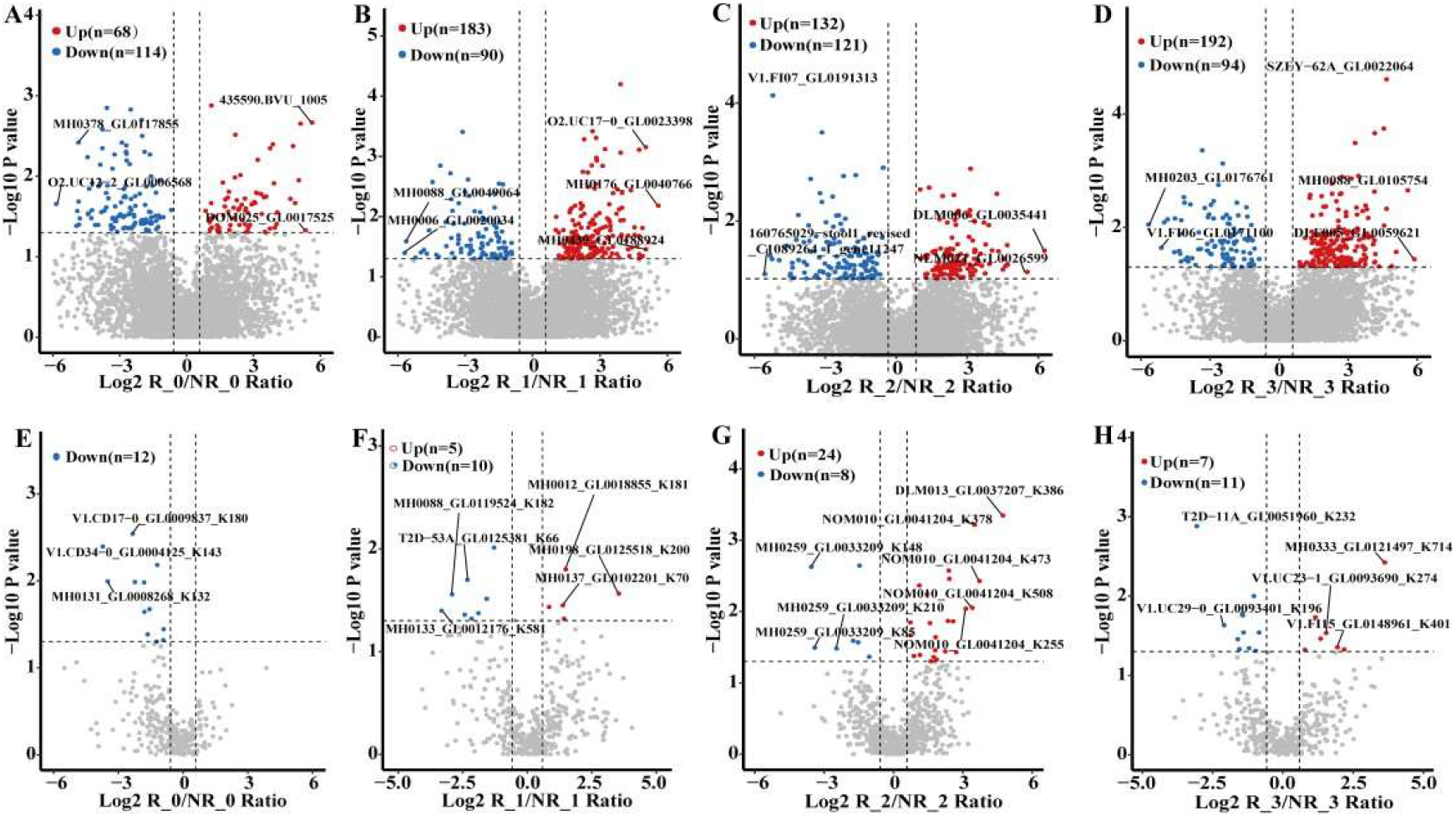
Differential protein abundance and Kcr sites between responders and non-responders across FMT time points in the microbiome. (A-D) Volcano plots of differential protein abundance at baseline (A), week 1 (B), week 8 (C), and week 12 (D). (E-H) Volcano plots of differential Kcr sites at corresponding time points. Significantly higher/lower features in responders are in red/blue.

We next classified differentially abundant features into baseline-altered and treatment-emergent signatures (Fig. 3). Eight proteins fulfilled the criteria for baseline-altered signatures, differing before FMT and maintaining a consistent direction of difference during follow-up, thereby representing candidate pretreatment markers of response (Figs. 3A, 3B). Treatment-emergent signatures were not different at baseline but became differentially abundant after FMT, reflecting response-dependent functional and post-translational remodeling (Figs. 3C-3F). Among these features, responders showed increased crotonylation of the beta subunit of CO dehydrogenase/acetyl-CoA synthase (CODH/ACS) and decreased crotonylation of 3-hydroxyacyl-CoA dehydrogenase (HADH), implicating acyl-CoA-centered metabolic pathways^11,12^ (Figs. 3C, 3D). Host inflammatory proteins, including S100A9 and serine protease inhibitors, showed reduced Kcr levels in responders after FMT^13-16^. In parallel, microbial proteins involved in SCFA production, nutrient transport, and polysaccharide degradation were upregulated in responders (Fig. 3E), supporting enhanced microbial metabolic restoration and ecological fitness.

**Fig. 3.**
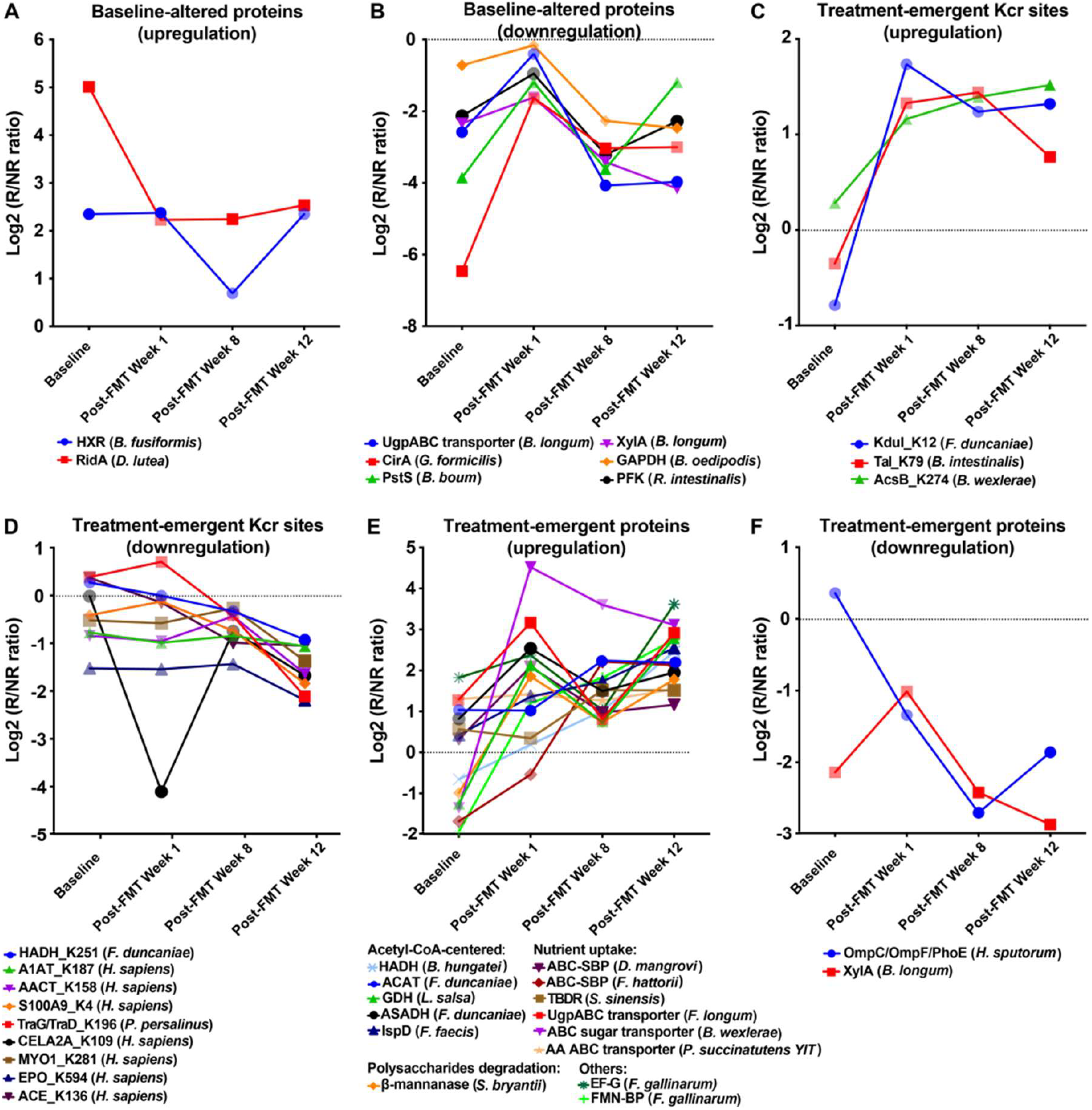
Temporal dynamics of baseline-altered and treatment-emergent molecular signatures. (A, B) Baseline-altered protein signatures. (C, D) Treatment-emergent Kcr signatures. (E, F) Treatment-emergent protein signatures. Line plots show longitudinal trajectories of individual features. Opaque symbols indicate significant differences. All abbreviations are defined in the Abbreviations section. A complete list of protein and Kcr signatures is provided in Table S1.

### FMT-responsive protein and Kcr features with group-specific and opposite regulation

To resolve the directionality of FMT-associated remodeling, we compared molecular changes from baseline to week 12 within each group and categorized features as responder-specific, non-responder-specific, or oppositely regulated (Fig. 4). Microbial Kcr sites on enzymes such as HADH and acetyl-CoA acetyltransferase showed response-specific regulation (Fig. 4A). A substantial set of host Kcr sites on inflammatory proteins, including S100A8/A9 and lactotransferrin, were specifically downregulated in responders (Fig. 4B), consistent with attenuation of neutrophil-associated inflammation. Microbial proteins involved in metal-ion transport and central metabolism were upregulated in responders but downregulated in non-responders (Fig. 4C). Conversely, host proteins related to neutrophil activation decreased in responders, whereas proteins linked to epithelial energy metabolism increased in non-responders (Fig. 4D), suggesting persistent inflammatory and metabolic stress in patients without clinical remission.

**Fig. 4.**
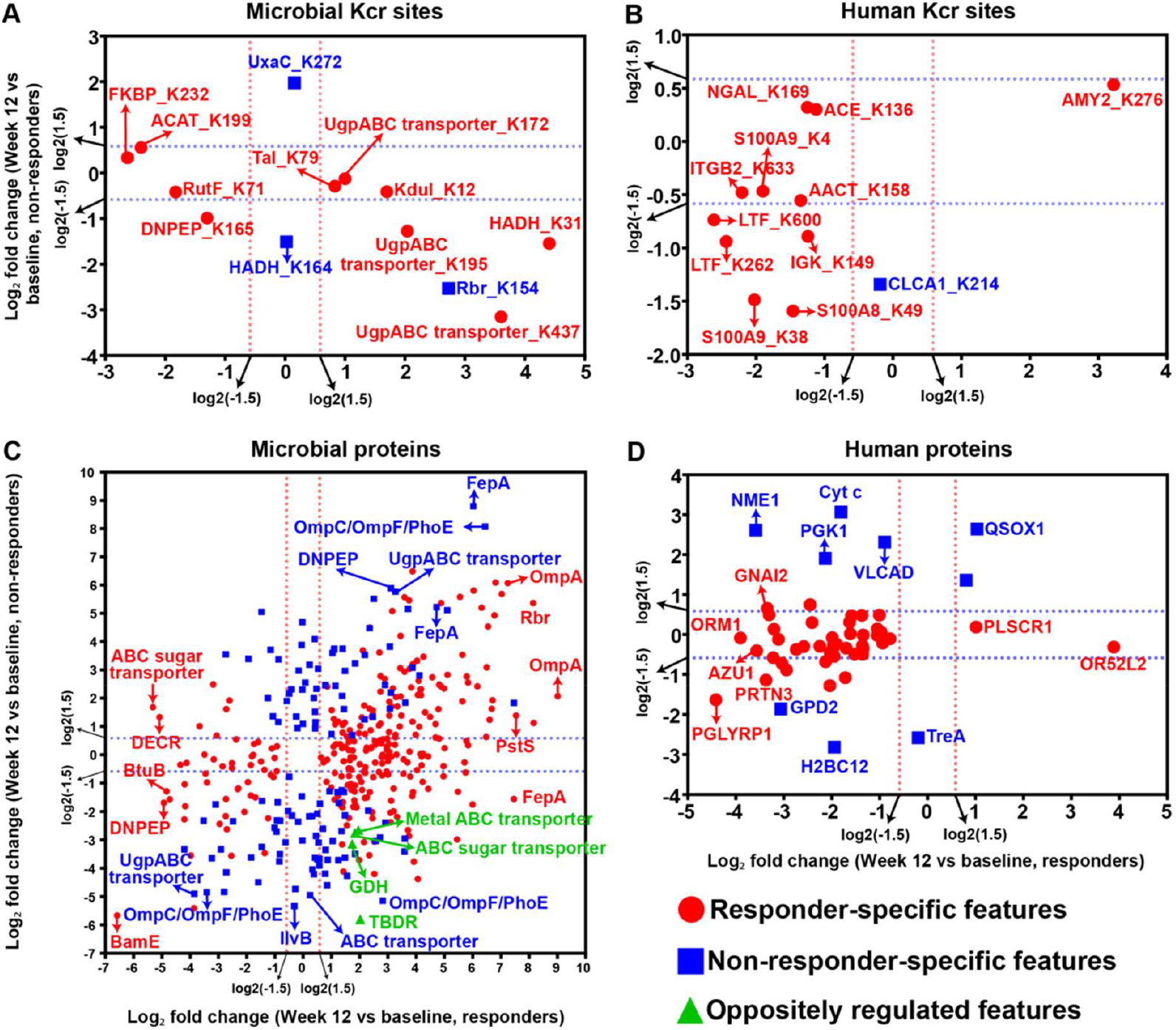
Group-specific and oppositely regulated molecular features post-FMT. Scatter plots of log2 fold change (week 12 vs. baseline) in responders (x-axis) vs. non-responders (y-axis) for (A) microbial Kcr sites, (B) host Kcr sites, (C) microbial proteins, and (D) host proteins. Points are colored by regulation category. All abbreviations are defined in the Abbreviations section.

## DISCUSSION

This study applies longitudinal fecal metaproteomics and lysine crotonylation profiling to a clinical UC-FMT cohort and identifies functional molecular features associated with treatment response. Rather than treating FMT efficacy as a taxonomic endpoint alone, our data indicate that clinical response is accompanied by coordinated remodeling of active microbial proteins, metabolite-sensitive Kcr sites, and host inflammatory signatures.

Four findings emerge. First, pretreatment molecular differences, including eight directionally consistent baseline-altered proteins, distinguish future responders from non-responders. Second, successful FMT is associated with increased microbial proteins and Kcr features related to SCFA and acyl-CoA-centered metabolism. Third, responders show coordinated attenuation of host inflammatory protein and Kcr signatures, particularly those linked to neutrophil activity. Fourth, non-responders show minimal or opposite remodeling, supporting the interpretation that clinical non-response reflects incomplete functional microbial restoration and persistent host inflammatory activation.

Our findings support lysine crotonylation as a dynamic and metabolite-sensitive post-translational layer during FMT response^11,12^. The concordance between microbial SCFA/acyl-CoA-related proteins, Kcr remodeling, and reduced host neutrophil-associated signatures provides a plausible molecular bridge between microbial metabolic restoration and mucosal inflammatory resolution. In this context, host inflammatory Kcr features involving proteins such as S100A8/A9 extend established fecal inflammatory markers and neutrophil-associated IBD biology into a post-translational framework for disease activity monitoring^13-16^.

Several limitations should be considered. The cohort was single-center and modest in size, and the baseline molecular signatures should be regarded as candidate pretreatment markers requiring validation in larger, independent FMT cohorts. Taxonomic inference was based on metaproteomic peptides rather than paired metagenomic sequencing, and the proposed SCFA/acyl-CoA-Kcr axis requires direct validation by targeted metabolomics, strain-level donor-recipient engraftment analysis, and mechanistic experiments. Future studies incorporating quantitative microbiome profiling, metabolomics, metagenomics, and rigorous confounder control will be important for distinguishing causal response mechanisms from response-associated molecular correlates^17-21^.

In conclusion, longitudinal metaproteomic and Kcr profiling provides a functional molecular view of FMT response in UC. The identified candidate pretreatment features and treatment-emergent remodeling patterns advance understanding of response heterogeneity and provide a foundation for future biomarker validation and mechanism-guided optimization of microbiota-targeted therapy.

## METHODS

### Sample collection

Patients with active UC were recruited from the IBD clinic at the First Affiliated Hospital of Kunming Medical University according to predefined inclusion and exclusion criteria. Sample size estimation was informed by publicly available metaproteomic data from a previously published UC cohort^22^. Differentially abundant proteins were used to construct a composite UC proteomic signature, and molecular restoration after FMT was modeled as a proportional reduction of the UC-healthy signature difference, supported by prior UC-FMT data showing post-treatment similarity between responders and donor microbiota^23^. Under an 80% restoration scenario, the corresponding planning effect size was estimated at d approximately 1.52, and G*Power analysis indicated that at least 9 participants per response group would provide 80% power at a two-sided alpha of 0.05. To account for biological variability, we aimed to enroll at least 10 participants per group. Accordingly, 23 patients with active UC were included, comprising responders (n = 13) and non-responders (n = 10). A total of 87 fecal samples were collected at baseline (responders, n = 11; non-responders, n = 10), week 1 (responders, n = 13; non-responders, n = 10), week 8 (responders, n = 13; non-responders, n = 10), and week 12 after FMT (responders, n = 12; non-responders, n = 8) (Table S2). All participants provided written informed consent, and the study was approved by the Institutional Ethical Review Board of the First Affiliated Hospital of Kunming Medical University. Fecal samples were collected in sterile tubes within 1 h of defecation and stored immediately at -80 °C until analysis.

### Donors for FMT

Healthy adults aged 18-45 years who had lived in Kunming for an extended period were screened using a standardized questionnaire^24^. Eligible candidates underwent stool and blood testing to exclude transmissible infections and multidrug-resistant organisms, including antimicrobial resistance (AMR) gene testing in fecal samples. Donor stool was processed using the FMT separation system (FMT-6A-50/12-AS, Nanjing Famaite) to obtain a fresh fecal suspension, which was administered to the ileal region through a colonoscopic tube within 30 min. Each administration used 200 mL of fecal suspension, and 600 mL was prepared from every 100 g of donor stool.

### Protein extraction and trypsin digestion

Frozen samples were homogenized in lysis buffer (1% Triton X-100, 1% protease inhibitor, 3 μM TSA, 50 mM NAM) and centrifuged (12,000g, 10 min, 4°C). Protein concentration was determined via BCA assay. Equal protein amounts were precipitated with 20% TCA, washed with cold acetone, and digested with trypsin (1:50, w/w) in 200 mM TEAB overnight. Reduction (5 mM DTT, 56°C, 30 min) and alkylation (11 mM IAA, RT, 15 min, dark) were performed before digestion.

### Peptide enrichment and LC-MS/MS analysis

Peptides were enriched using antibody-conjugated resin (PTM0503, PTM Bio) in IP buffer (100 mM NaCl, 1 mM EDTA, 50 mM Tris-HCl, 0.5% NP-40, pH 8.0) with overnight incubation (4°C). After washing, bound peptides were eluted with 0.1% TFA, desalted (C18 ZipTips), and vacuum-dried. LC separation was performed on a Nano Elute system with a gradient of 7-80% mobile phase B (0.1% FA in acetonitrile) at 450 nL/min. Eluted peptides were ionized (1.65 kV) and analyzed by timsTOF Pro 2 in PASEF mode (MS/MS range: 100-1700 m/z, dynamic exclusion: 24 s).

### Metaproteomics and meta-crotonylome analysis

Metaproteomic analysis used an iterative database search strategy adapted from MetaPro-IQ^25,26^. Raw MS data were first subjected to spectral clustering with PRIDE Cluster to reduce redundancy^27^. Clustered spectra were searched against a comprehensive protein database using X! Tandem with trypsin/P specificity^28,29^. Carbamidomethylation of cysteine was set as a fixed modification, while methionine oxidation and protein N-terminal acetylation were set as variable modifications; lysine crotonylation was included as an additional variable modification for Kcr datasets. Up to four missed cleavages were allowed, and protein identification was controlled at 1% FDR. Identified protein sequences were used to construct integrated microbiome protein databases containing 262,492 entries for the metaproteome and 126,504 entries for the meta-crotonylome. MS/MS spectra were then searched with MaxQuant (v1.6.15.0)^30^ against the integrated microbiota database concatenated with the UniProt human proteome database (20,429 entries; accessed 20 December 2023). Target-decoy FDR thresholds were set at 1% for both proteins and peptides. Kcr peptide levels were quantified using raw spectral intensity and normalized to the corresponding protein abundance to reduce confounding by protein-level changes.

### Taxonomic annotation

All unique microbial peptides were submitted to Unipept (v2.2.1) for microbial taxonomic analysis using the lowest common ancestor (LCA) algorithm, with equal I/L and advanced missed-cleavage handling enabled. Peptides that could not be assigned to a specific taxonomic level due to conservation across multiple domains of life were classified as root. Relative taxon abundance was normalized to the fraction of all peptides at each taxonomic rank. Quantified microbial proteins were annotated with COG categories using EggNOG 5.0 and EggNOG-mapper with default parameters.

### Identification of differentially abundant proteins and Kcr sites between responders and non-responders

To identify protein and Kcr features that longitudinally distinguish FMT responders from non-responders, we compared their abundance between responders and non-responders at baseline and at post-treatment time points. For each time point, statistical significance of differences between groups was assessed using the Mann-Whitney U test. P values were adjusted using the Benjamini-Hochberg procedure. Features with an FDR < 0.05 and an absolute fold change >= log2(1.5) were considered differentially abundant. This approach was applied consistently to both protein-level and Kcr peptide-level data.

### Classification of baseline-altered and treatment-emergent signatures

Differentially abundant protein and Kcr features identified across time points were categorized according to their temporal patterns. Baseline-altered signatures were defined as features that differed significantly between responders and non-responders at baseline and maintained a consistent direction of difference throughout post-treatment time points. Treatment-emergent signatures were defined as features with no significant difference at baseline that became differentially abundant between groups after FMT. This classification separated candidate pretreatment features from response-dependent molecular changes during treatment.

### Identification of FMT-responsive Kcr peptides and proteins with group-specific and opposite regulation

To investigate molecular changes associated with FMT efficacy, we evaluated alterations in the abundance of Kcr sites and proteins from baseline to 12 weeks after FMT separately in responders and non-responders. For each group, within-group comparisons were performed to identify features that changed significantly following treatment. Statistical significance was assessed using the Mann-Whitney U test, followed by Benjamini-Hochberg correction. Features with an FDR < 0.05 and an absolute fold change >= log2(1.5) were considered significantly altered. Features were classified a responder-specific, non-responder-specific, or oppositely regulated according to direction and significance across groups.

### Statistical analysis

Continuous variables were expressed as mean ± SD or median depending on distribution, and categorical variables were expressed as percentages. Continuous variables were compared with the Mann-Whitney U test, and categorical variables were compared with the χ2 test or Fisher’s exact test.

## ABBREVIATIONS

A1AT: Alpha-1-antitrypsin
AA ABC transporter: ABC-type amino acid transport system
AACT: Alpha-1-antichymotrypsin
ABC-SBP: ATP-binding cassette transporter substrate-binding protein
ABC sugar transporter: ABC-type sugar transport system, periplasmic component
ABC transporter: ATP-binding cassette transport system
ACAT: Acetyl-CoA acetyltransferase
ACE: Angiotensin-converting enzyme
AcsB: CO dehydrogenase/acetyl-CoA synthase beta subunit
AMR: Antimicrobial resistant
ASADH: Aspartate-semialdehyde dehydrogenase
AZU1: Azurocidin
B. boum: Bifidobacterium boum
B. fusiformis: Blautia fusiformis
B. hungatei: Butyrivibrio hungatei
B. intestinalis: Blautia intestinalis
B. longum: Bifidobacterium longum
B. oedipodis: Bifidobacterium oedipodis
B. wexlerae: Blautia wexlerae
BamE: Outer membrane protein assembly factor BamE
BtuB: Outer membrane cobalamin receptor protein
CELA2A: Chymotrypsin-like elastase family member 2A
CirA: Colicin import membrane protein
CLCA1: Calcium-activated chloride channel regulator 1
CODH/ACS: CO dehydrogenase/acetyl-CoA synthase
Cyt c: Cytochrome c
D. lutea: Daejeonella lutea
D. mangrove: Donghicola mangrove
DECR: NADPH-dependent 2,4-dienoyl-CoA reductase
DNPEP: Aspartyl aminopeptidase
EF-G: Translation elongation factor G
EPO: Eosinophil peroxidase
FMT: Fecal microbiota transplantation
FDR: False discovery rate
F. duncaniae: Faecalibacterium duncaniae
F. faecis: Faecalibacterium faecis
F. gallinarum: Faecalibacterium gallinarum
F. hattorii: Faecalibacterium hattorii
F. longum: Faecalibacterium longum
FepA: Outer membrane ferrienterochelin receptor
FKBP: FK506-binding protein-type peptidyl-prolyl cis-trans isomerase
FMN-BP: FMN-binding domain-containing protein
GAPDH: Glyceraldehyde-3-phosphate dehydrogenase
GDH: Glutamate dehydrogenase/leucine dehydrogenase
GNAI2: G protein subunit alpha i2
GPD2: Glycerol-3-phosphate dehydrogenase, mitochondrial
G. formicilis: Gemmiger formicilis
H. sapiens: Homo sapiens
H. sputorum: Haemophilus sputorum
H2BC12: Histone H2B type 1-L
HADH: 3-Hydroxyacyl-CoA dehydrogenase
HXR: Hydroxylamine reductase
IBD: Inflammatory bowel disease
IGK: Immunoglobulin kappa light chain
IlvB: Acetolactate synthase large subunit
IspD: 2-C-methyl-D-erythritol 4-phosphate cytidylyltransferase
ITGB2: Integrin beta-2
Kcr: Lysine crotonylation
KduI: 5-Keto-4-deoxyuronate isomerase
L. salsa: Litchfieldia salsa
LTF: Lactotransferrin
MYO1: Myosin-1
NGAL: Neutrophil gelatinase-associated lipocalin
NME1: Nucleoside diphosphate kinase A
NR: Non-responder
OmpA: Outer membrane protein A
OmpC/OmpF/PhoE: Outer membrane porins C, F, and E
OR52L2: Olfactory receptor 52L2
ORM1: Alpha-1-acid glycoprotein 1
P. persalinus: Pseudocohnilembus persalinus
P. succinatutens YIT: Phascolarctobacterium succinatutens YIT
PFK: 6-Phosphofructokinase
PGK1: Phosphoglycerate kinase 1
PGLYRP1: Peptidoglycan recognition protein 1
PLSCR1: Phospholipid scramblase 1
PRTN3: Proteinase 3 (myeloblastin)
PstS: ABC-type phosphate transport system, periplasmic component
PTM: Post-translational modification
QSOX1: Quiescin sulfhydryl oxidase 1
R: Responder
R. intestinalis: Ruminococcus intestinalis
Rbr: Rubrerythrin
RidA: Enamine deaminase RidA
RutF: FMN reductase RutF
S100A8: Protein S100-A8
S100A9: Protein S100-A9
SCFA: short-chain fatty acid
S. bryantii: Segatella bryantii
S. sinensis: Segatella sinensis
Tal: Transaldolase
TBDR: TonB-dependent receptor
TraG/TraD: Transfer protein G/D family TPase
TreA: Trehalase
UC: Ulcerative colitis
UgpABC transporter: ABC-type glycerol-3-phosphate transport system
UxaC: Glucuronate isomerase
VLCAD: Very long-chain acyl-CoA dehydrogenase
XylA: Xylose isomerase
β-mannanase: Endo-1,4-β-mannanase.

## DECLARATIONS

### Ethics approval and consent to participate

This study was approved by the Ethics Committee of the First Affiliated Hospital of Kunming Medical University (Approval No. (2025) Lun Shen L No. 111) and was conducted in strict accordance with the principles of the Declaration of Helsinki, CFDA/GCP guidelines, and relevant laws and regulations of China. The ethical committee’s responsibilities, composition, operational procedures, and records comply with ICH-GCP standards. All participants provided written informed consent (Version 1.0, dated March 13, 2025) after being fully informed about the study’s purpose, procedures, potential risks, and benefits. The informed consent form was reviewed and approved by the Ethics Committee.

### Patient and Public Involvement Statement

Patients and the public were involved in the design, conduct, reporting, and dissemination plans of this research. Specifically, patient representatives were consulted on the study design and outcome measures to ensure clinical relevance. Feedback from patient advisory input was incorporated into the interpretation of results and preparation of the manuscript.

### Availability of data and material

The in-house metaproteomic and lysine crotonylation raw data, along with the corresponding normalized abundance profiles, have been deposited in the National Omics Data Encyclopedia (NODE; https://www.biosino.org/node/) under accession number NODE: OEP00006925.

### Authors’ contributions

Y.S., W.L., and Y.M. conceived and designed the study. Y.L., Y.Y., X.L., and B.L. conducted sample collection and related experiments. W.L. and W.C. performed omics data processing and analysis. W.L. and Y.Y. drafted the manuscript, and all authors contributed to its revision. R.Z. and L.Z. provided senior supervision and guidance.

## Acknowledgments

We acknowledge the computational support provided by the Center for Scientific Computing and the Supercomputing Center of the School of Life Sciences and Technology, Tongji University.

